# Vertical transmission of core endophytes through the seeds

**DOI:** 10.1101/2025.01.06.628327

**Authors:** Irene Sanz-Puente, Santiago Redondo-Salvo, Gloria Torres-Cortés, María de Toro, Susana Fernandes, Andreas Börner, Óscar Lorenzo, Fernando de la Cruz, Marta Robledo

## Abstract

Plant-associated microorganisms, particularly endophytes, are essential for plant health and development. Endophytic microbiota is intimately associated with host plants colonizing various tissues, including seeds. Seed endophytes are particularly noteworthy because of their potential for vertical transmission. This pathway may play a role in the long-term establishment and evolution of stable bacteria-host interactions across plant generations. Hundreds of seed-bacteria associations have been recently uncovered; however, most seem to be transient or unspecific. While it is known that microorganisms can be transmitted from plant tissues to seeds and from seeds to seedlings, the experimental confirmation of bacterial transfer through successive plant generations remains unreported. In this study, we identified *Pantoea* as the unique core endophytic bacteria inhabiting the endosperms of 24 wheat seed samples originally harvested in different worldwide locations. Remarkably, *Pantoea* is the genus with the highest relative average abundance in wheat seeds (61%) and also in germinated seedlings grown under gnotobiotic conditions (30%). In the field, it was the only genus dwelling roots, shoots, spikes and seeds of 4 different wheat varieties tested and its abundance progressively increased across these tissues. This genuine pattern of vertical enrichment, which was not found in other common wheat-associated taxa, suggests a role in the transfer of these endophytic bacteria through the seeds. To confirm intergenerational transmission, parental plants were inoculated with labelled *Pantoea* isolates, which specifically colonized the next generations of Poaceae plants, experimentally demonstrating bacterial vertical inheritance to the offspring generations and suggesting transmission specificity.

## INTRODUCTION

All multicellular organisms host complex microbial communities [1]. In humans, the maternal microbiome helps establish progeny intestinal flora, immune system and metabolism [2]. Plants also host diverse microbial communities, forming the “holobiont” [3]. A healthy plant holobiont, mainly prokaryotes, fosters plant-microbial homeostasis [4,5]. Microorganisms that colonize internal plant tissues without causing apparent damage are critical for plant health [6,7], including symbiotic endophytes [6]. One well studied example is the nitrogen-fixing bacteria *Rhizobium leguminosarum,* which can survive in the soil or internally colonize legume root nodules to fix nitrogen [8]. Interestingly, certain rhizobial strains can also colonize cereal or vegetables as growth-promoting endophytes [9,10].

Research traditionally focused on rhizosphere and phyllosphere communities [3]. However, the microbiota associated with reproductive organs, which may be potentially involved in vertical transmission, is recently receiving more attention [11–14]. Seeds, essential for plant regeneration, were long considered axenic [15,16]. However, seed tissues harbor complex microbial communities, which may exert beneficial or deleterious effects on plant growth and health [4,17]. In this research direction, recent studies have identified the presence of bacteria in internal seed structures [18,19].

Three major transmission pathways of seed-borne microorganisms have been suggested: external (via seed surface contact), floral (through the stigma), and internal (via the vascular system) [12,17,20]. The external pathway is considered the most permissive route, while internal transmission is probably more restricted to endophytes [12,21]. Microscopic studies have recently confirmed the presence of vertically-transmitted *Burkholderia* in the flower buds, close to the embryos, but not in the vascular tissues [22]. However, the pathogen *Xanthomonas* was observed in connections of maternal vascular tissues to seeds and in the embryo [23,24]. Microbial transmission from seeds to seedlings is well studied [25], particularly in relation to pathogens [26,27]. Amplicon sequencing has suggested the transmission of certain endophytic taxa across plant generations [28–31]. Several works isolating the same bacterial species from both G0 and G1 seeds also indicate natural transmission [28,32,33]. A study carrying out microbiota sequencing analysis of seeds over several generations even suggests that only few endophytes might be consistently transmitted [28]. However, overlaps among microbiota members across tissues is not sufficient to prove vertical transmission and strain-specific identification is required to confirm seed-to-seed transmission of certain bacterial taxa.

Confirming vertical transmission is thus technically challenging. Several studies have verified that labelled GPF- or GUS bacteria provide a reliable approach for tracking endophytes seed dynamics [23]. Fruit and flower inoculation of GUS-labelled *Paraburkholderia phytofirmans* PsJN, isolated from onion, demonstrated endophytic colonization of the next generation in grapevines and maize [34,35]. *Arabidopsis* roots inoculated with GFP-labelled *Bacillus thuringiensis* yielded bacteria in seedlings [36]. However, to the best of our knowledge, vertical transmission of native seed endophytes through plant generations has not been experimentally confirmed.

Considering that vertical transmission is more likely to be the widespread phenomenon among ubiquitous bacteria dwelling in the endosperm, this work aimed to identify core endophytes across different varieties of wheat (*Triticum sp.*), one of the most cultivated cereals worldwide [37]. We characterized the endophytic microbiota of commercial and ancestral wheat samples from different global origins. Sequencing analysis revealed that wheat seed endophytic microbiota is dominated by *Pantoea*. This dominance is maintained upon seedling germination in axenic conditions, but wild plants exhibit a *Pantoea* gradient across tissues, finally accumulating again in seeds. To investigate vertical inheritance, we set up a 3-year field experiment and established a model system based on GUS-labelled bacteria that enabled us to document seed bacterial plant colonization across three plant generations.

## MATERIALS AND METHODS

### Plant material and soil sampling

The endophytic bacterial community of wheat was studied using 24 seed samples from two main sources (Table S1-S3): plant material collected from fields in Spain at the end of the growing season and seeds harvested worldwide and regenerated at IPK germplasm bank, (Gatersleben, Germany). Rhizospheric soil surrounding wheat roots was also sampled in sterile tubes. At the laboratory, plant samples were divided into root, shoot, spike, and mature seeds harvested from spikes. Samples were stored in paper bags at 4°C before processing. For greenhouse experiments, seeds from Tae_SP4 (Table S3), *Lolium multiflorum*, and *Arabidopsis thaliana* Col-0 were used.

### Plant surface disinfection and DNA extraction

Plant tissues (∼0.5 g) were placed in 1 mL PBS (Phosphate Buffered Saline with 0.05% Tween-20) and sonicated for 1’ (Ultrasons, Barcelona, Spain). Three surface disinfection methods were tested [7,24,34], and the latter was routinely used. Briefly, seeds, spikes, shoots, and roots were rinsed in 70% ethanol 3’ (5’ for roots), followed by 5% Active chlorine for 10’ (5’ for seeds). Samples were then washed three times with sterile distilled water, and 100 µL of the final wash were plated on TSA (Tryptic Soy Agar, Condalab, Madrid, Spain) at 30°C for 3 days to confirm disinfection.

To isolate endophytic bacteria, surface-disinfected material (∼0.25 g) was disrupted and soaked in 1 mL PBS overnight at 4°C. Serial dilutions were plated on TSA and incubated at 30°C. The DNeasy PowerLyser PowerSoil Kit (Quiagen, Hilden, Germany) was utilized for bacterial DNA extraction from plant tissues following manufacturer’s instructions. DNA was eluted in 50 µl purified water and quantified using Nanodrop (ThermoFisher, Massachusetts, United States).

### Microbial communities sequencing and analysis

The V4 region of the 16S rRNA gene was amplified using primers 515_F and 808_R with Illumina adapters (Table S4). PCR reactions contained 0.5 µM of each primer, 200 µM dNTPs, 0.02 U/µL Kapa2G Robust, and 25 ng of the corresponding DNA. PCR included an initial denaturation 98°C (30’’), 25 cycles (95°C, 15’’; 55°C, 15’’; 72°C, 10’’), and a final extension (72°C, 5’). Amplicons were purified with Agencourt^®^ AMPure^®^ XP and sent to the CIBIR Genomics Core Facility (La Rioja, Spain) for qualification (Fragment Analyzer, Agilent) and quantification (Qubit HS DNA Kit, ThermoFisher). Barcodes (Nextera XT, Illumina) were added before sequencing on an Illumina MiSeq (PE300).

Sequencing provider-delivered paired FASTQ files underwent quality assessment with FastQC (v0.11.9), trimming with Cutadapt (v1.18), and denoising with DADA2 (v1.6.0) to generate Amplicon Sequence Variants (ASVs). Taxonomic assignment was performed using Qiime2 (v2019.1) against the SILVA v132 database. Non-bacterial sequences and ASVs with <0.1% abundance were excluded to reduce diversity overestimation [38]. Raw data can be found in the supplemental information.

Ecological analyses were conducted in R (v2023.06.1). Phyloseq objects facilitated data integration, taxonomic profiles α- and β-diversity were computed with Vegan, and visualized with ggplot2.

### Whole Genome Sequencing (WGS) of *Pantoea* seed isolates

Consecutive generations of *T. aestivum var. Chambo* seeds were harvested from a field in León (Spain) and bacterial endophytes were isolates as described before. Yellow colonies were identified via full 16S rRNA gene sequencing (27_F, 1522_R, and 515_F; Table S4) using Sanger sequencing. Whole genomes of *Pantoea* isolates were sequenced by MicrobesNG (Birmingham, United Kingdom) using Illumina short-read technology. DNA extraction, sequencing, quality control, demultiplexing, and trimming of raw reads were performed by the provider. Trimmed reads were assembled into contigs with Unicycler (v0.5.1). SNPs among *Pantoea* genomes were identified using Snippy (v4.6.0).

### GUS Labelling of Pantoea agglomerans

Plasmid pGUS-3 [39] was conjugated into *P. agglomerans* C-88, isolated from wheat seeds, via biparental mating with *E. coli* S17-1 (λpir). The conjugation mixture was plated on TSA with kanamycin (50 µg/mL) and incubated overnight at 28°C. Transconjugants were transferred to TSA plates containing the chromogenic substrate X-gluc (1 µL/mL, Biogen). Degradation of X-gluc by β-glucuronidase encoded by pGUS-3 resulted in blue colonies after overnight incubation at 28°C. The presence of the plasmid was confirmed by PCR using GUS_F and GUS_R primers (Table S4), yielding strain *P. agglomerans* C88- GUS.

### Gnotobiotic plant germination and growth

Disinfected seeds were placed on 1% agar plates with 1 mL sterile distilled water and incubated in the dark at 4°C 48 hours (*Arabidopsis*) or overnight (Tae_Sp2) for stratification. All plates were then incubated at 24°C in the dark for 48 hours to allow germination. Seedlings were individually transferred to sterilized glass tubes containing 20 mL Murashige and Skoog Basal salt mixture (Sigma:M5519) and a filter paper. Tubes were sealed with cotton, covered and placed in a growth chamber (16/8 hour light/dark, 24°C, 60% humidity). After 7 days, shoots and roots were excised under sterile conditions for DNA extraction and amplicon sequencing.

### Plant growth conditions and bacterial inoculation

*P. agglomerans* C88-GUS overnight cultures on TSA were suspended in PBS at an OD_600nm_ of 0.6 (approximately 4x10^8^ CFUs/mL). G0 surface-disinfected seeds of wheat, *Lolium*, and *Arabidopsis* were germinated as described. Seedlings were transferred to pots with a 50% soil-vermiculite mixture and grown in a greenhouse (16/8 hour light/dark photoperiod, 18-30°C, 60% relative humidity) until flowering.

For bacterial incorporation into seeds (Figure S3B), *Arabidopsis* flowers were individually sprayed with 50 µL of the C88-GUS suspension, followed by removal of the floral bud. For *Lolium* and wheat, spikes were immersed in 40 mL of the same suspension. To assess bacteria transmission across three generations (Figure S3C), G0 seeds were imbibed in a C88-GUS suspension for 24 hours, sown into pots, and grown in the greenhouse until flowering. Seeds from the subsequent G1 generation were harvested and stored at 4°C until further analysis. All control plants were treated with PBS without bacteria.

### Detection of *P. agglomerans* C88-GUS in plant tissues

G1 and G2 seeds were surface-disinfected and germinated as previously described. Seven-day-old seedlings were immersed in GUS buffer containing 50 mM phosphate buffer, 0.05% Triton X-100, 1 mM K_3_Fe(CN)_6_, 1 mM K_4_Fe(CN)_6_, 0.05 M EDTA, and 1.2 mM of X-gluc. Plants were incubated at 37°C in the dark until blue coloration appeared, washed twice and stored in 70% ethanol until imaging. Stained tissues were observed under a stereoscopic microscope (Olympus-SZX16/SZ12) and photographed with a NIKON-DS-Fi1 camera.

To confirm and quantify the presence of C88-GUS, seven-day-old surface-disinfected and germinated seedlings were split into shoot, root, and seed tissues. Triplets of pooled tissues were processed for DNA extraction as described before. The *gusA* gene, encoding β-glucuronidase, was amplified and quantified on a StepOnePlus™ Real-Time PCR System (ThermoFisher). Each 20 µl reaction contained 4 µl of PerfeCTa qPCRToughMix, 0.6 µl of primers (100 nM), and a TaqMan probe targeting the *gusA* gene (Table S4). PCR amplification included denaturation at 95°C (3’), followed by 40 cycles of 95°C (15’’), 60°C (30’’), and 72°C (30’’). Fluorescence data were collected, and Ct values were determined using StepOnePlus software. Relative quantification was based on a standard curve correlating CFUs/g of C88-GUS mock-inoculates in seedlings with Ct values. Negative controls without DNA were included in all reactions.

## RESULTS

### *Pantoea* is present in the core microbiota of all wheat seed varieties

To assess bacterial endophytic communities, first we set a reliable protocol to facilitate wheat seed disinfection. We removed epiphytic microorganisms and soil particles by seed sonication, followed by the surface-disinfection method described by Mitter et al (2017), ensuring total removal of bacterial seed external load (Figure S1). Next, to identify core seed endophytes, we harvested 12 wheat samples in Spain (Table S1) and studied their endophytic bacterial communities by 16S rRNA sequencing. The wheat species analyzed were categorized according to their genome content into commercial and ancestral species (Table S1) based on their approximate dates of divergence (before and after 0.13 million years ago, respectively) [40]. Our metabarcoding analysis showed that wheat seed endophytic bacterial assemblages are relatively conserved at genus level disregarding this categorization, with a predominant presence of *Pantoea* (Figure 1A).

**Figure 1.**
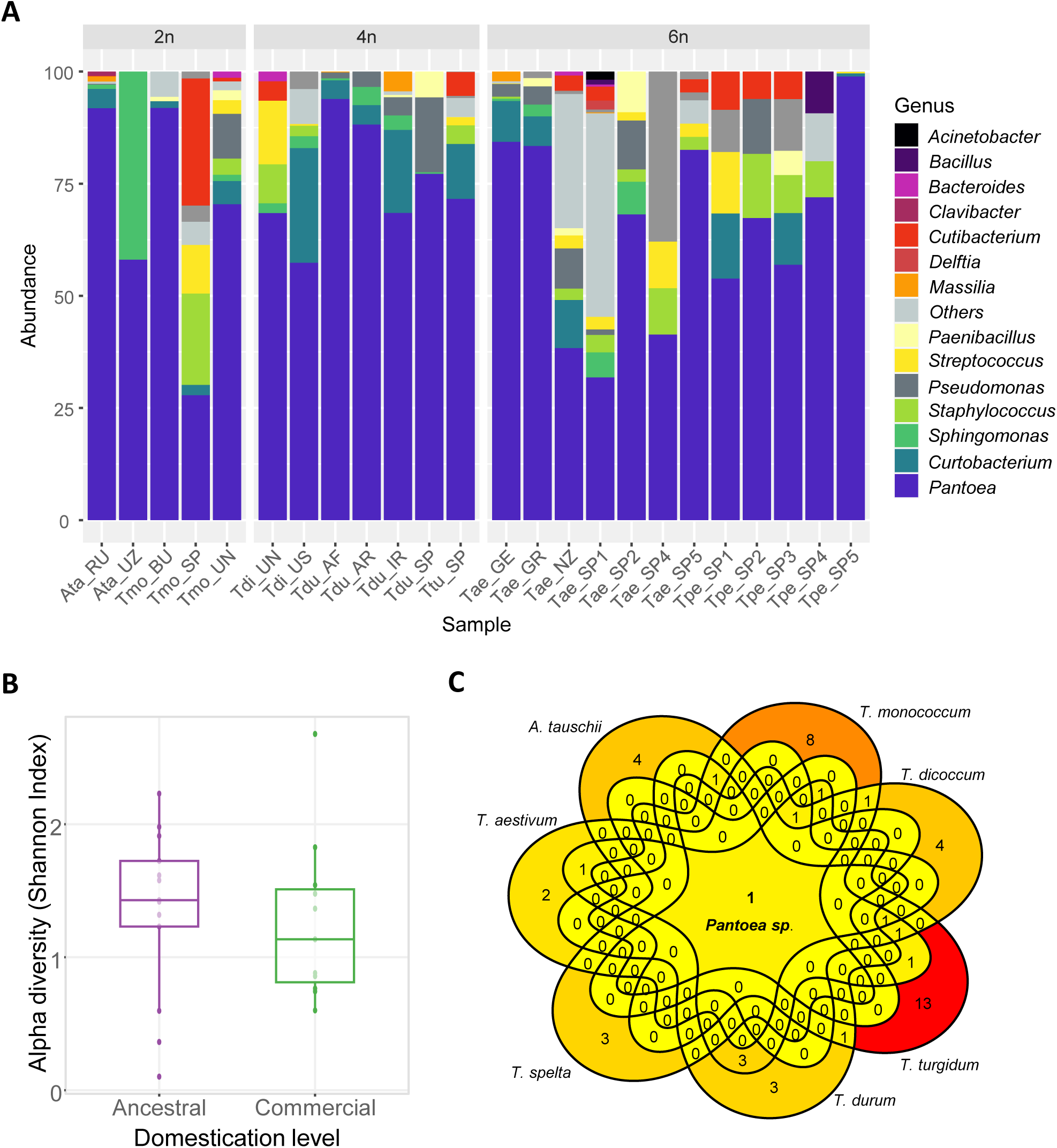
Endophytic bacterial diversity of worldwide wheat seeds. A) Relative abundance of dominant bacterial genus (>2%) present in wheat seeds from the diploid (2n) species *Aegilops tauschii* (Ata) and *Triticum monococcum* (Tmo), as well as tetraploid (4n) *T.turgidum* (Ttu), *T. dicoccum* (Tdi) and *T. durum* (Tdu); together with hexaploid (6n) *T. aestivum* (Tae) and *T. spelta* (Tpe; Tables S1 & S2). B) α-diversity metrics (Shannon index) of samples grouped according to plant domestication level. Each box represents the Shannon index interquartile range and the line inside is the median. Whiskers extend up to the last sample in 1.5 times the interquartile range. C) Venn diagram illustrating the unique and common bacterial ASVs among the seven wheat-related seed species included in this work. Only shared bacterial ASVs present in all seed pools of the same wheat species are included. The areas of overlap represent elements common to two or more sets, while the non-overlapping sections show exclusive elements. Colors range from yellow to red (minimum to maximum number of ASVs).

To exclude a bias in sample location, we extended our set with 12 additional samples from the IPK seedbank originally collected on 4 different continents (Table S2). Surprisingly, the overall taxonomic composition was similar (Figure 1A). Besides *Pantoea*, with a 61% average abundance in the 24 wheat seed samples, the most common bacterial genus found were *Curtobacterium* (5%), *Staphylococcus*, *Sphingomonas* and other *Erwiniaceae* (3%). Measures of α-diversity showed an overall low diversity of seed endophytic bacterial features compared to other plant tissues (see below). Plant domestication, wheat species, genome content or original harvest location did not affect significantly bacterial abundance (Figures 1B, and S2A-C). The PCoA plot revealed no clear clustering or patterns in relation to these sample characteristics (Figures S2D-E), further supporting conservation of the seed endophytic assemblages among wheat varieties worldwide.

We next focused on the prevalence among bacterial communities inhabiting seeds of the seven wheat-related species included in this work. For that, all shared bacterial ASVs among seeds of the same wheat species were extracted to establish the core bacterial microbiome for each sample. To facilitate the identification of commonalities, the list of core seed bacterial ASVs for each wheat species was used to create a Venn diagram (Figure 1C). A total of 50 ASVs were analysed, revealing both unique and shared bacterial ASVs across the species. Common core bacterial ASVs among wheat species are scarce, featuring 12 ASVs shared by *T. durum* and *T. monococcum*. However, all seeds contain a single ubiquitous core endophytic genus present in all wheat samples from different worldwide locations. This unique bacterial genus has been taxonomically assigned to *Pantoea sp*.

### *Pantoea* is the most abundant genus in gnotobiotically germinated wheat seedlings

We next determined the seed community dynamics after germination under gnotobiotic conditions using a hydroponic cultivation system. The microbial communities of plant roots and shoots from seedlings obtained by metabarcoding were compared with those from non-germinated seeds. Analysis of α-diversity revealed that the detectable bacterial community in seedlings was significantly higher than in seeds (Figure 2A). Furthermore, β- diversity analysis (Figure 2B) indicated that seed data cluster separately from the post-germination communities. PERMANOVA significantly differed across tissues (R² = 0.81, p = 0.01), explaining 82.8% of the variation. Relative abundances of bacterial taxa in seeds and germinated roots and shoot are compared in Figure 2C. Most genera that are present in the seeds at low percentages proliferated upon germination in roots or/and shoots (e.g. *Curtobacterium*). Interestingly, *Pantoea* was the only ASV whose average relative abundance in seeds (73.3% in these 3 sample sets) decreased in germinated roots and shoots (30%). Nevertheless, *Pantoea* remains the most relative abundant bacterial member in both wheat roots and shoots 7 days after germination.

**Figure 2.**
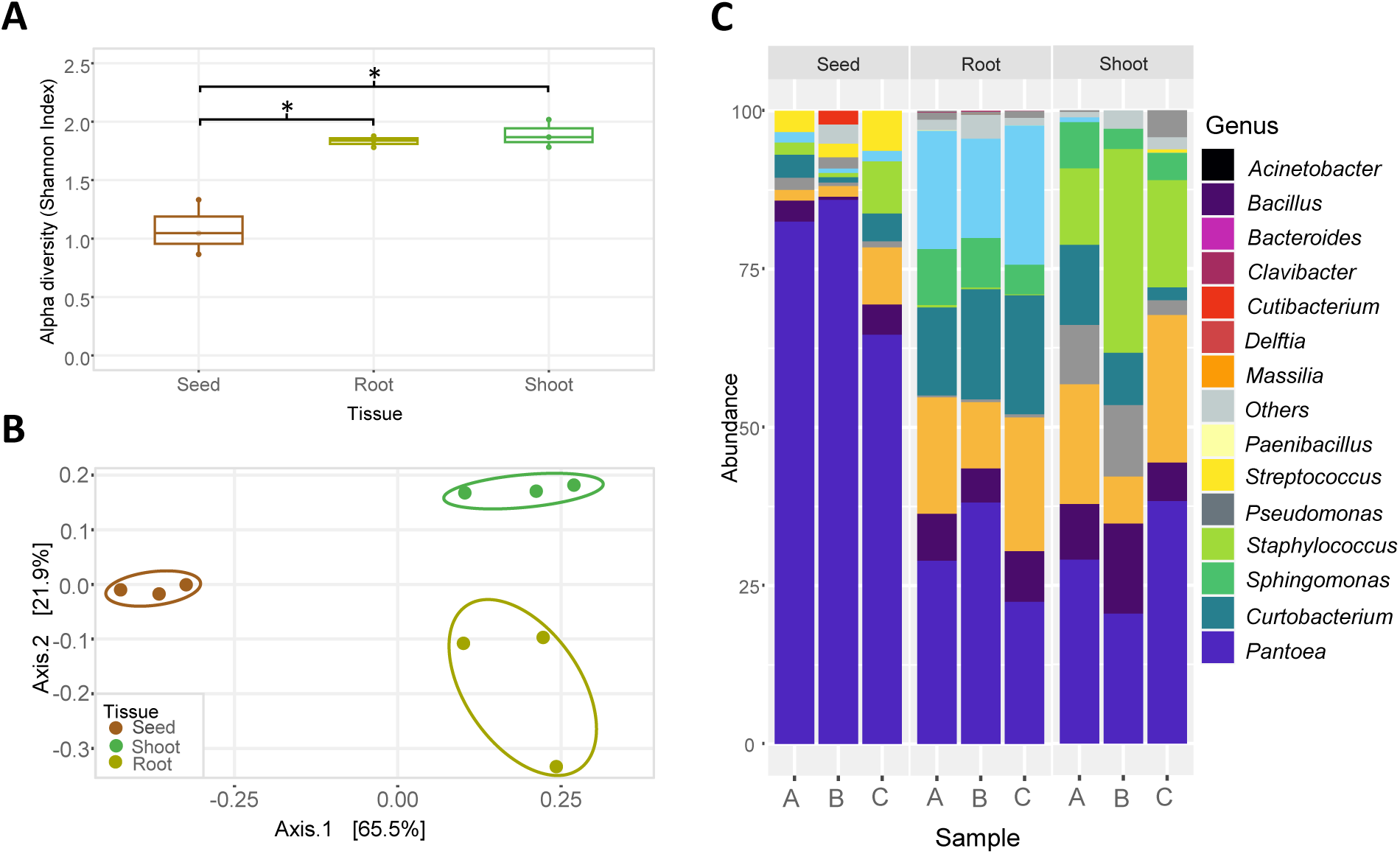
Endophytic bacterial diversity of wheat seedlings cultured under gnotobiotic conditions. A) Bacterial α-diversity (Shannon index) of Tae_SP2 surface disinfected seeds and gnotobiotically germinated shoot and roots. * p<0.05 (Tukey’s post-hoc test). B) PCoA corresponding to the Bray–Curtis dissimilarity index (β-diversity) of the bacterial communities present in the different plant tissues. Each dot corresponds to an individual technical replicate. The x- and y-axes represent the first and second components of the PCoA plot, respectively. Ellipses show the clustering and relative homogeneity of bacterial communities within tissues (PERMANOVA: R² = 0.81, p = 0.01). C) Bar plot of bacterial endophytic taxa present at >2% relative abundance in wheat seeds and gnotobiotically germinated seedlings. Three individual replicates are shown (A, B, C), corresponding to three sets of 12 surfaced-disinfected seeds from the same wheat variety sample (Tae_SP2).

### All wheat tissues grown under field conditions dwell *Pantoea*

Next, we analysed core endophytic bacterial profiles in wheat grown in field conditions. With this objective, we compared the communities inhabiting seeds with those present in different surface-disinfected plant tissues of four wheat species grown in the field (Table S3). The rhizospheric soil of each plant sample was also analysed. The endophytic community diversity of wheat plants grown under field conditions (Figure 3) was notably different from the obtained in gnotobiotic system (Figure 2). α-diversity analysis revealed significant differences among soil and all plant-derived communities across the wheat tissues compared (Figure 3A). The seed community exhibited the lowest diversity, which increased progressively in the order of seed < spike < shoot < root < soil. β-diversity analysis (Figure 3B) revealed that bacterial communities significantly differed across plant tissues (R² = 0.38, p = 0.001), explaining 38.39% of the total variation. However, no apparent pattern was shown when we attempted to statistically ordinate bacterial communities present in our samples by their plant genotype (R² = 0.15, p = 0.137). In accordance with the α- and β- diversity analyses, the obtained taxa bar plot showed that phylogenetic diversity gradually decreased from soils to seeds (Figure 3C).

**Figure 3.**
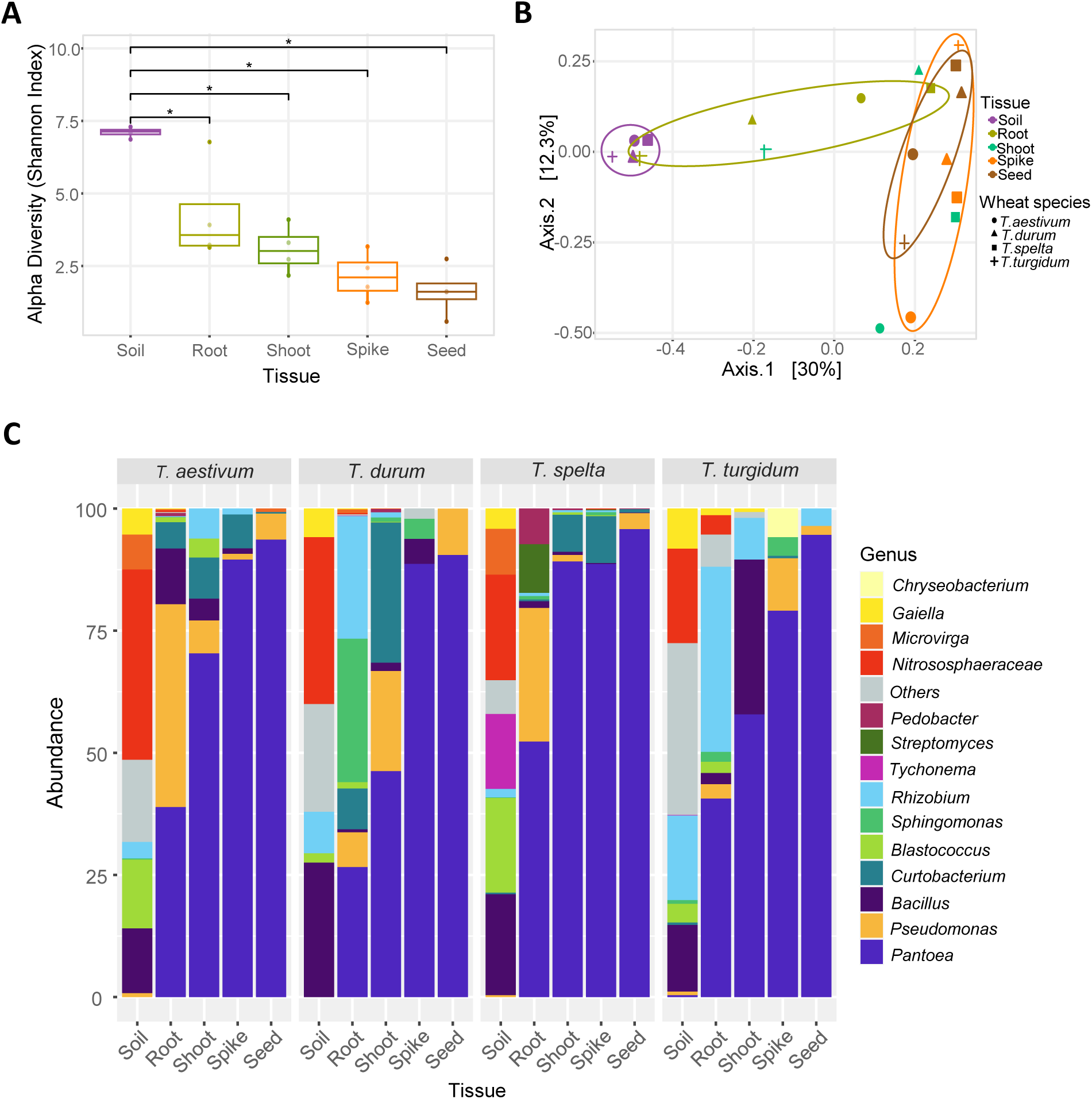
Endophytic bacterial diversity of wheat tissues and species grown under field conditions. A) Bacterial α-diversity (Shannon index) different wheat plant tissues and the surrounding soil. * p<0.05 (Tukey’s post-hoc test). B) β-diversity (PCoA) of the bacterial communities associated with different plant tissues and wheat species as indicated in the legend. Each dot corresponds to an individual biological replicate. Ellipses represent the clustering and relative homogeneity of bacterial communities within tissues (PERMANOVA: R² = 0.38, p = 0.001). C) Bar plot showing relative abundance of bacterial endophytic taxa present in >5% ASVs in soil and plant tissues of *Triticum aestivum*, *T. durum*, *T. spelta* and *T. turgidum*.

To establish the core bacterial microbiomes from this dataset, shared ASVs among all plant tissues of the same wheat species (Figure 4A) and among each plant tissue of different wheat species (Figure 4B) were extracted. Considering its notably difference (Figure 3), the soil community was excluded from this analysis. Most core ASVs present in each wheat sample are unique of the plant species, especially in *T. aestivum*. Curiously, only 4 bacterial ASVs (0.4%) are shared between the four wheat species: *Pantoea*, *Streptococcus, Pedobacter* and *Pseudomonas* (Figure 4A). Shared bacterial endophytes across the four wheat species comprised 14, 132, 64 and 927 ASVs in seeds, spikes, shoots, and roots, respectively (Figure 4B). Interestingly, only *Pantoea*, *Staphylococcus* and *Curtobacterium* were found in all wheat tissues (Figure 4B). Specifically, *Pantoea* is the only endophytic genus accompanying all wheat varieties and tissues in the field (Figures 4A and 4B).

**Figure 4.**
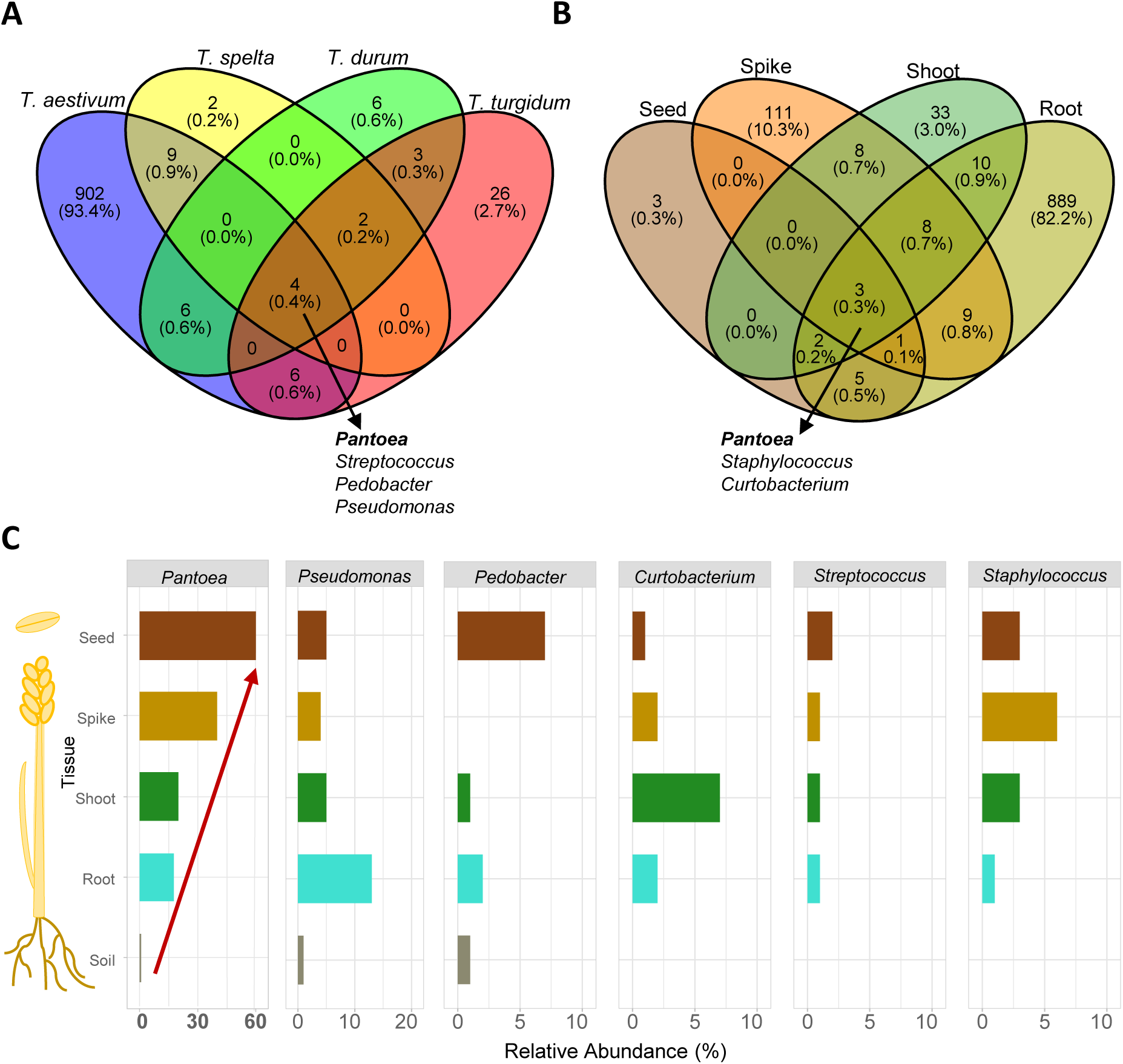
Wheat core endophytic bacteria dynamics during plant field development. Venn diagrams illustrating unique and shared bacterial ASVs, along with their respective proportions, among *Triticum* species (A) and tissues (B). Taxonomical assignment of core ASVs present in all samples is indicated. C) Average relative abundance in plant tissues of core endophytic genera found in all analysed plant tissues or all wheat samples (pannels A and B). Note that *Pantoea* abundance in the field gradually rises from root to seed wheat tissues (red arrow).

### *Pantoea* abundance shows a distinct increase pattern from root to seed wheat tissues

Next, we studied wheat core bacteria dynamics during plant development by comparing the abundance of the shared populations among the four wheat species or tissues. Population dynamics of *Pantoea,* which was not detected in most soil samples, exhibited the lowest abundance in roots and increased progressively in the order of root < shoot < spike < seed in all analysed wheat species (Figure 3C). This vertical gradient was particularly pronounced in *T. durum* and *T. spelta*, where relative abundances of *Pantoea* increased sequentially from root to seed, ranging from 23-38-58-67% and 24-33-56-84%, respectively. Seeds were the most preferentially colonized by this genus, disregarding the wheat species (Figure 3C).

We next aimed to compare bacterial-tissue enrichment patterns of the 6 identified core ASVs in Figures 4A-B. For that, we calculated their average coverage among wheat plant species in each plant tissue and the soil (Figure 4C). Based on this, we confirmed again the *Pantoea* clear vertical enrichment from roots to seeds. Curiously, other genus like *Pseudomonas* mostly preferred roots for colonization, while *Curtobacterium* accumulated in shoots and *Staphylococcus* in spikes (Figure 4C). However, *Pantoea* distinct pattern is not mirrored in none of the other core plant-associated taxa. Therefore, this is the only core taxa conserved across host genotypes and tissues displaying an abundance vertical enrichment gradient (Figure 4C, red arrow).

### Vertical transmission of *Pantoea* to the offspring plant generation through the seeds

The consistent presence and abundance of *Pantoea* in the seeds, along with its vertical enrichment throughout plant tissues, suggests potential transmission through the seeds to subsequent generations. To test this hypothesis, we first grew *T. aestivum* seeds over 3 generations in the field (Figure S3A). Seed endophytic microbiota of each generation (G0 to G2) was plated and *Pantoea* isolates were identified by 16S rRNA gene sequencing. Most isolates belonged to *P. agglomerans*. WGS of three *P. agglomerans* isolates belonging to the three generations (PG0 to PG2) revealed a high degree of similarity among them. Specifically, a single SNP distinguished a *P. agglomerans* strain isolated from the first wheat generation (C-113) from another isolated in the third generation (C-204).

Isolation of nearly identical *Pantoea* strains across generations strongly supports the hypothesis of vertical transmission through wheat seeds. To experimentally confirm this theory, we labelled a *Pantoea* wheat seed isolate and tracked it across plant generations. For that, we selected the strain *P. agglomerans* C-88, isolated from surface disinfected *T. spelta* seeds (Tpe_SP4). This strain was stably transformed with a plasmid encoding β- glucuronidase, leading to C88*-*GUS. In a first set of greenhouse experiments depicted in Figure S3B, *T. aestivum* spikes were inoculated with a GUS-labelled bacterial suspension as detailed in the methods section. Seeds of the following progeny were germinated and stained to detect GUS activity. As shown in Figure S4A, 7-day-old wheat seedlings derived from spikes inoculated with *Pantoea* C88*-*GUS show a clear blue staining, in contrast to roots from uninoculated control samples. *Pantoea* quantification in these G1 germinated seedlings was performed through qPCR by interpolating Ct values in a standard curve (Figure S4B). Prior to DNA extraction, G1 seedlings were split into shoot, root and the remaining seed. Matching the colorimetric results, *Pantoea* C88*-*GUS preferably colonized G1 wheat roots, reaching 1.4 x 10^4^ CFUs/g of tissue. To investigate if *Pantoea* vertical transmission ability was conserved in a plant phylogenetic framework, we performed the same experiment with seeds from the forage grass *Lolium multiflorum* (Poaceae) and the model plant *Arabidopsis thaliana* (Brassicaceae). Curiously, we confirmed *Pantoea* incorporation and transmission in *Lolium* but not in *Arabidopsis* (Figure S4A).

Both qualitative and quantitative methods support that *Pantoea* can be incorporated to the seeds of wheat and transmitted to the germinating seedlings. Finally, to confirm bacterial vertical inheritance from seeds to seedlings of the G2 progeny we set up a new set of experiments (Figure S3C). *L. multiflorum* and *T. aestivum* seeds were imbibed with the labelled endophyte and germinated seedlings were transferred to plant pots until flowering. After harvest, the next generation of seeds were germinated again, and seedlings were stained with X-gluc to verify the presence of the endophyte as described before (Figure 5A). Interestingly, the results obtained by GUS histochemical analysis and qPCR methods matched, showing that C88-GUS significantly accumulated in the root seedlings (Figure 5B). Most importantly, these results confirm that *Pantoea* C88-GUS can be vertically inherited to the offspring generation through the seeds in both Poaceae.

**Figure 5.**
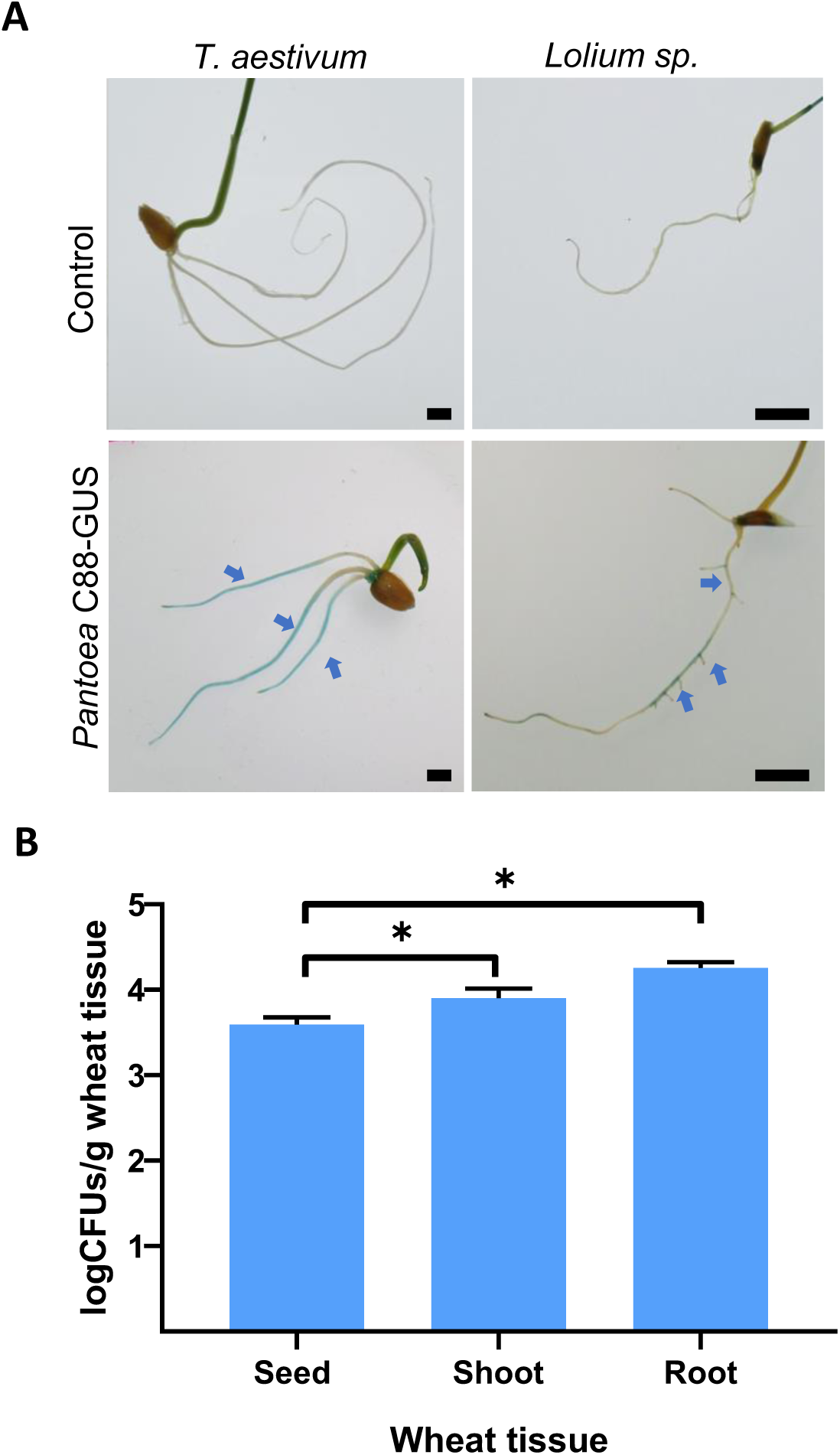
Experimental confirmation of *Pantoea* C88-GUS vertical transmission from inoculated G0 seeds to G2 seedlings. A) *Pantoea* colonization of G2 seedlings assessed by X-gluc histochemical analysis. Blue arrows indicate the presence of the endophyte *Pantoea* C88-GUS in *T. aestivum* and *L. multiflorm* (Poaceae). B) Quantification of *Pantoea* population in wheat tissues via qPCR. Error bars represent the standard deviation of the mean from three independent experiments with three biological replicates per experiment. Bar 5 mm.

## DISCUSSION

Seed microbial reservoir is involved in bacterial transmission, as inferred from shared species in seeds and seedlings [28–31]. However, the presence of common species does not necessarily implicate vertical transmission, as considerable heterogeneity exists within taxa from the same species. Experimental confirmation is needed to verify steady vertical transmission of plant-associated bacteria. This study demonstrates bacterial vertical transmission events of a bacterial endophyte through plant generations providing novel insights into the acquisition and community dynamics of the seed microbiome.

We hypothesized that core endophytic seed bacteria widespread among a worldwide- cultivated cereal like wheat will be ideal candidates for confirming vertical transmission. Most studies aiming to widely characterize the seed-associated microbiota do not perform a profound surface-disinfection of the endosperms [41]. The role and diversity of bacteria inhabiting seed endosphere is yet under-explored, mainly due to technical difficulties and the absence of standardized methodologies [24,42]. Therefore, we firstly aimed to establish a robust surface disinfection protocol to eliminate the abundant seed epiphytes. Our experiments provide evidence that sonication and Mitter et al. (2017) protocol led to reliable seed surface-disinfection, removing epiphytic bacteria. These is one of the first attempts to provide a validated framework for studying seed endophytes and is applicable across other plants.

As corroborated by previous literature, seeds harbor low-diverse endophytic bacterial communities, mostly dominated by Gamma-Proteobacteria and Firmicutes [25,43,44]. α- and β-diversity analyses suggest that harvest location, wheat species, genome content, or domestication level do not majorly influence bacterial diversity, supporting the absence of a rational taxonomical structure as suggested before [27]. Previous studies also showed no significant differences between commercial and ancestral wheat seed-derived communities [44], while others reported higher diversity among cultivated cereals [45]. However, the differences in plant varieties, seed disinfection and sequencing methods hamper reliable comparisons. In agreement with our results, a recent meta-analysis of 63 seed microbiota studies reported that *Pantoea* was one of the most abundant and prevalent seed-borne genera, being present in 27 different plant species [46]. This suggests that these members of the endophytic communities may be essential for development and adaptation of different plant species. *Pantoea,* belonging to Erwiniaceae, is a diverse facultatively anaerobic genus of yellow-pigmented bacteria. *Pantoea* comprises over 20 recognized species that show a remarkable ecological adaptability, being frequently isolated from diverse environments, especially in association with plants [47,48]. While some species, like *P. stewartii,* show pathogenic traits [49]; other like *P. agglomerans* possess plant growth-promoting and biocontrol abilities [47]. Curiously, this species was the most frequently isolated in our wheat seed samples.

Microbial succession during germination differs between plant species [24,25,41,50]. Our gnotobiotic system confirms that not all wheat seed endophytic taxa are necessarily transmitted to seedlings [25,43]. Interestingly, in the absence of bacteria from the rhizosphere, the dominance of *Pantoea* is maintained in plant roots and shoots upon germination. In contrast, we noticed an increase in bacterial phylogenetic diversity in wheat roots and shoots grown under field conditions, indicating the replacement of this dominant pioneer seed endophytic taxa by soil-derived microorganisms. *Pantoea* persisted across wheat tissues from all wheat species that we scrutinized, and its relative abundance gradually increases from seeds to roots. Our results also indicated that relative abundances of other common bacterial genera did not show a *Pantoea*-like consistent gradient across wheat tissues, suggesting that microbial community assembly is tissue-specific. Nevertheless, this intriguing accumulation pattern may not be necessary for vertical transmission. Environmental drivers and host pressures shaping wheat microbiota warrant further study [51].

*Pantoea* unique vertical enrichment through wheat plant tissues and the isolation of *P. agglomerans* strains with almost identical genomes across three wheat generations supported transfer of these endophytic bacteria through the seeds. To experimentally confirm intergenerational transmission endophytes, we tracked the wheat seed isolate *P. agglomerans* C-88 colonization ability across plant generations. Our results confirm that bacterial endophytes can be vertically transferred via seeds to the progeny through plant generations. Interestingly, GUS-tagged isolates were detected in *T. aestivum* and *L. multiflorum* (Poaceae), but not in *A. thaliana* (Brassicaceae), suggesting host-specificity in vertical transmission. This hypothesis is consistent with previous studies showing that seed-associated microbiota exhibits host specificity, which may be driven by co- evolutionary dynamics, selective pressures imposed by the host plant immune system or distinct bacterial transmission pathways [27,32]. *P. agglomerans* was also one of the three bacterial OTUs detected in all radishes (Brassicaceae) unsterilized seed samples across sequencing of three successive plant generations [29]. Therefore, it is tempting to speculate that different strains of *P. agglomerans* have evolved the ability to be vertically transmitted across generations in Brassicaceae and other plant families, perhaps as part of a conserved trait among seed-transmitted symbiotic endophytes [6,21].

Interestingly, *Pantoea* was not detected in most wheat rhizospheric soil samples, further supporting its genuine vertical transmission through wheat seeds. This finding aligns with the theory that vertically-transmitted endophytes may form stable, co-evolved relationships with their host plants [6]. This symbiosis-like association may be shaped by selective pressures that favor non-pathogenic strains [52] together with the benefits gained by both the bacteria and the plant. The microbial counterpart ensures its dispersion through the seeds, survival, and niche displacement of microbial competitors while the host plant benefits by providing their progeny with beneficial symbionts [20]. Plant microbiome functions like germination are essential for some hosts [53]. Additionally, cosmopolitan plant-associated bacteria, including *Pantoea, Stenotrophomonas, Bacillus* and *Pseudomonas*, positively impact germination [54]. In particular, *P. agglomerans* PS1 significantly increased wheat seed germination up to 25% [55]. Altogether, these findings suggest that *Pantoea* may establish a beneficial symbiotic relationship favouring both germination and plant development.

The fact that *Pantoea* was the only genus identified in all our analyzed wheat seed samples agrees with the taxonomical restriction of microorganisms that seem to consistently pass on to progeny plants [29,56]. This further suggests that the ability to be transmitted from seed to seed across plant generations is not a widespread trait among bacteria and that there must be strict filtering processes governing these mechanisms. Similarly, some microorganisms are directly transferred from mother to baby during birth, although few persist from birth to adulthood in the offspring [57]. Investigation into these mechanisms, probably related to plant defense from seed-derived pathogens, deserves further studies.

Interestingly, *Pantoea* ability to be transmitted to the progeny has also been well-documented in insects [58]. Some Hemiptera establish obligate symbiotic associations with a wide diversity of *Pantoea,* which are vertically inherited from adult females to nymphs. To our knowledge, only the leaf-nodulating nitrogen-fixing *Burkholderia* symbionts were described as obligate in plants [59,60]. However, *Burkholderia*-free host plants survived in a sterile *in vitro* environment [22], suggesting that even for obligate symbionts vertical transmission may not be the only route for plant survival. The ability of surface sterilised wheat embryos to germinate leading to axenic seedlings [7], suggests the absence of obligate endophytes also in these cereal plants, at least under laboratory conditions.

Our findings suggest that transmission from parent to progeny is a common strategy among *Pantoea* endophytes, but the genetic and ecological factors driving host-specificity require further research [6]. Vertical transmission of endophytes persisting in the seeds over several host plant generations may enable a tight interaction or even plant-endophyte coevolution. The conservation and dynamics of this microbial genus suggests a potential evolutionary symbiosis-like relationship maintained between *Pantoea* and wheat throughout its domestication process. Our results confirm that vertical transfer of bacterial endophytes occurs, but whether obligate and strictly vertically transferred symbioses with bacteria is a widespread phenomenon or plants usually acquire new microbiota with each generation remains obscure. In particular, the role of environmental filtering and host selection in shaping microbial transmission routes warrants further investigation. Future studies exploring the co-evolution of plants and their microbial partners in early developmental stages could provide valuable insight into the long-term stability and functional significance of these symbiotic interactions.

## Acknowledgements

This work was funded by the research grants to FdlC./M.R.: PID2020-117923GB-I00, CPP2022L009595 funded by MCIN/AEI/10.13039/501100011033 and the “European Union” or by the “European Union NextGenerationEU/PRTR”; and Spanish Centre for Technological Development and Innovation (CDTI) grant (IDI20200826C). M.R. has been granted PTQ-17-09029 and RYC2022-035122-I funded by MCIN/AEI. I.S-P. was granted by CVE:2019-8472 from Cantabria government. We would like to thank the Leibniz Institute of Plant Genetics and Crop Plant Research (IPK), Gatersleben Germany, for providing the seed material, Matthieu Barret for kindly revising the manuscript and all the members of the Intergenomics group for stimulating discussions.

## Author contributions

M.R., G. T-C. and F.dlC. contributed to the conception and design of the work; I.S-P., S.R. M.dT., S.F., and M.R. performed the experimental assays; I.S-P., S.R. and M.R. analyzed the data, I.S-P. and M.R. drafted the manuscript; and A. B., O. L. and F.dlC. substantively revised it.

## Data Availability Statement

Raw reads from WGS are available in the NCBI GenBank repository under the accession number PRJNA1201053. Metabarcoding raw data generated in this study have been included as a supplement to this publication.

---

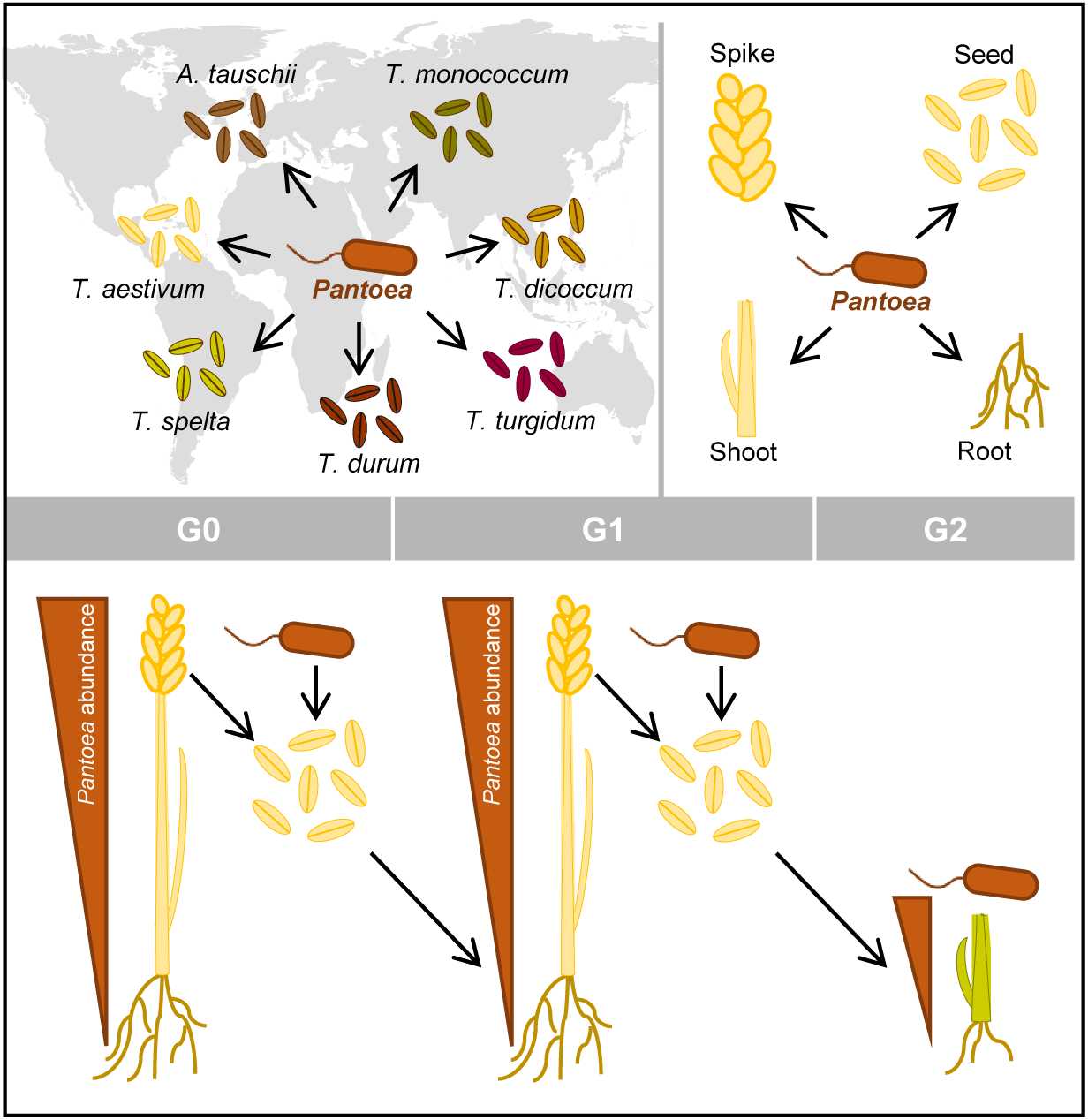

