## Supplemental Material for "Vertical transmission of core endophytes through the seeds"

This document contains supplemental figures and tables for the indicated manuscript

Total number of pages of Supporting Information: 9 (including cover page)

Number of Figures in Supporting Information: 4

Number of Tables in Supporting Information: 4

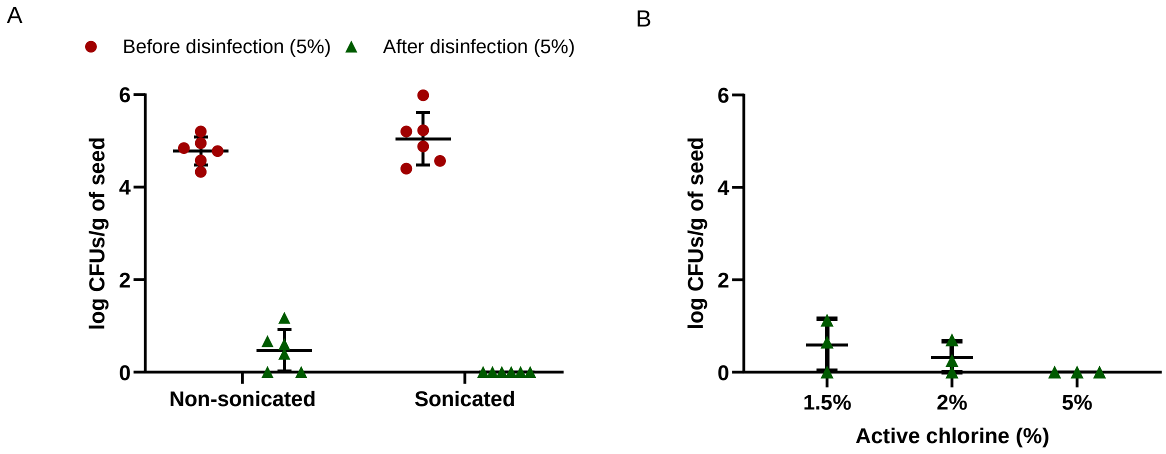


**Figure S1. Optimization of wheat seed endophytic microbiome analysis.** A) Bacterial load in non-sonicated and sonicated pooled samples of wheat (*Triticum aestivum* var Craklin) seeds. Colony Forming Units (CFUs) were measured before (red points) or after (green points) surface disinfection with 5% active chlorine. Each point represents a technical replicate from each of the two independent assays (each biological replicate is a pool of 12 seeds). Error bars indicate SD. B) Comparison of three wheat seed disinfection methods, presented as CFUs per seed gram after disinfection with three different active chloride concentrations: 1.5% (Robinson et al., 2016); 2% (Torres-Cortés et al., 2018); and 5% (adapted from Mitter et al., 2017). Each point represents a measure from three independent assays (with 3 pools of 12 seeds per replicate). Error bars indicate SD.

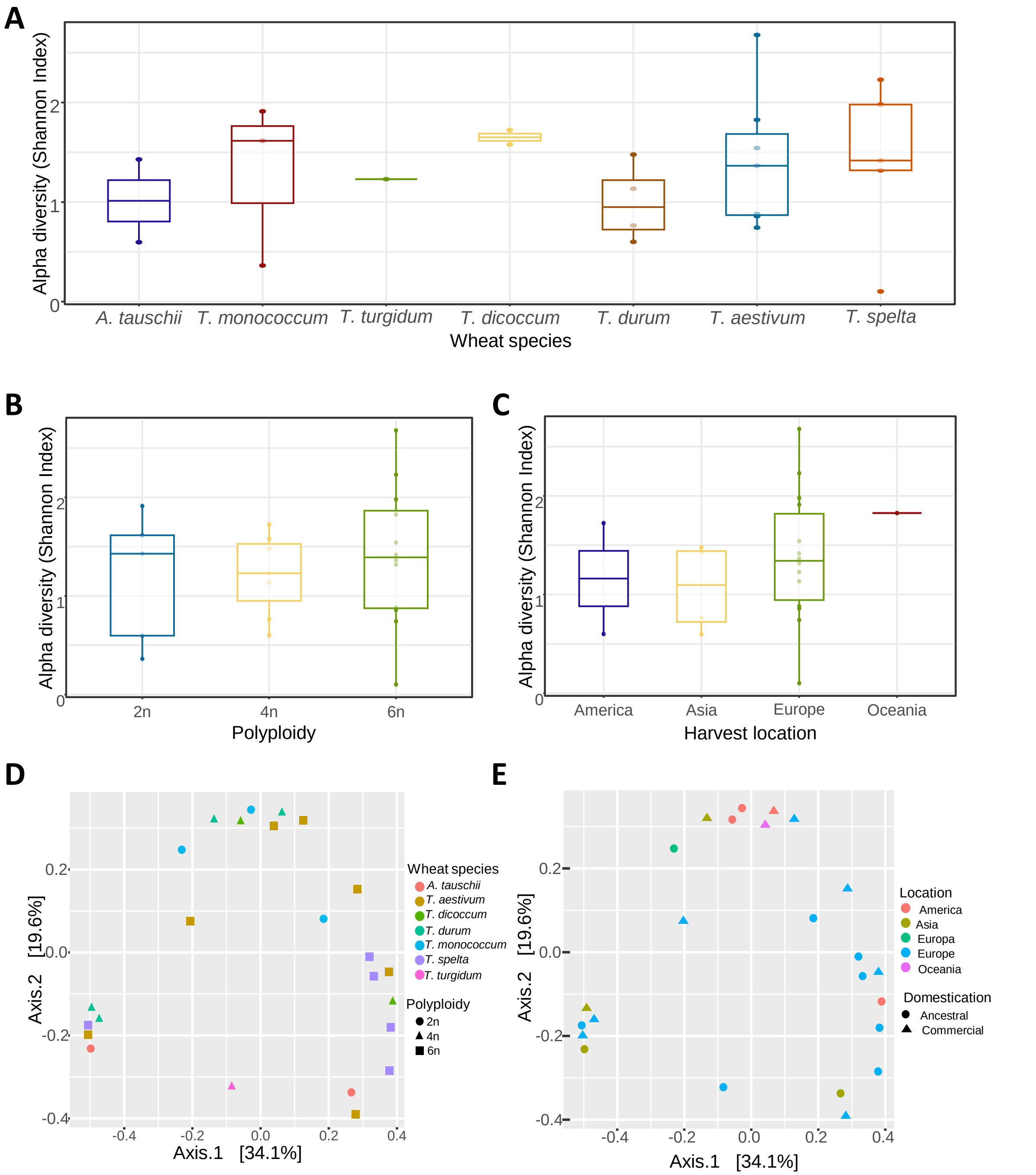


**Figure S2. Analysis of wheat seed bacterial worldwide diversity.** Box-and-whisker plots show α-diversity metrics (Shannon index) of samples grouped according to wheat species (A), polyploidy (B) or harvest location (C). Each box represents interquartile range and the line inside the box is the median. D, E) PCoA corresponding to the Bray–Curtis dissimilarity index (β-diversity) of the bacterial communities present in wheat seeds depicted according to the legends. Each dot represents an individual technical replicate. The x- and y-axes represent the first and second components of the PCoA plot, respectively. PERMANOVA revealed that bacterial seed communities are similar no matter the host wheat species, polyploidy, harvest location or domestication ($R^2$ = 0. 2).

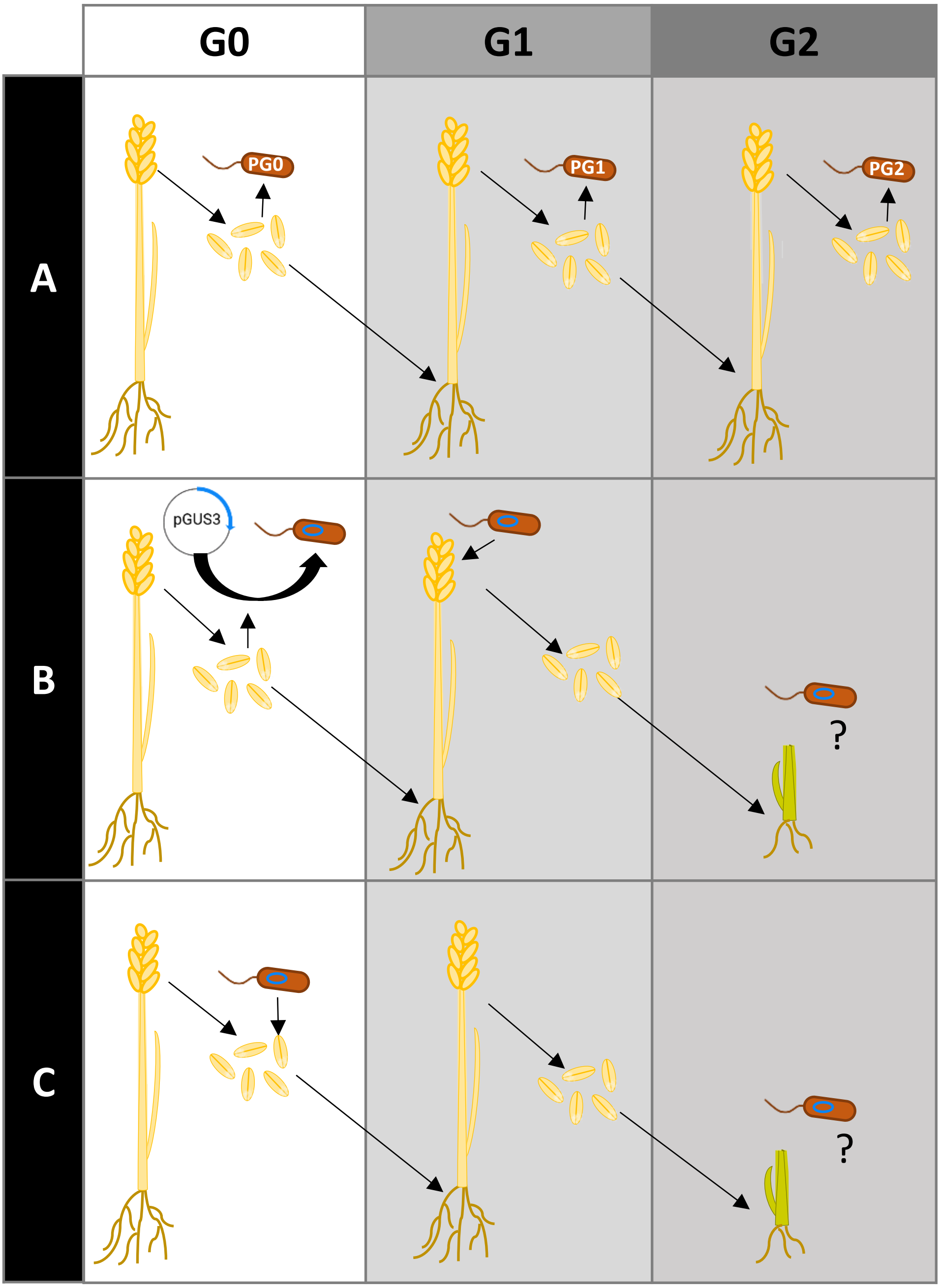


**Figure S3. Schemes of the different experimental setups to assess *Pantoea* vertical transmission across consecutive wheat generations**. A) The diagram illustrates three successive generations of wheat plants grown in the field (G0, G1, and G2). *Pantoea* strains (PG0, PG1, PG2) were isolated from seeds belonging to each plant generation for genome sequencing. B) These panels described the first set of greenhouse experiments: plant flowers were inoculated with a GUS-labelled *P. agglomerans* seed isolate and seeds of the G2 progeny were germinated and stained to detect bacterial colonization. Results are shown in Figure S4. C) In the second set of experiments seeds were imbibed with the labelled endophyte and grown until flowering. The next generation of seeds were germinated, and seedlings were stained to verify the presence of the endophyte. Results are shown in Figure 5. See details in the main text.

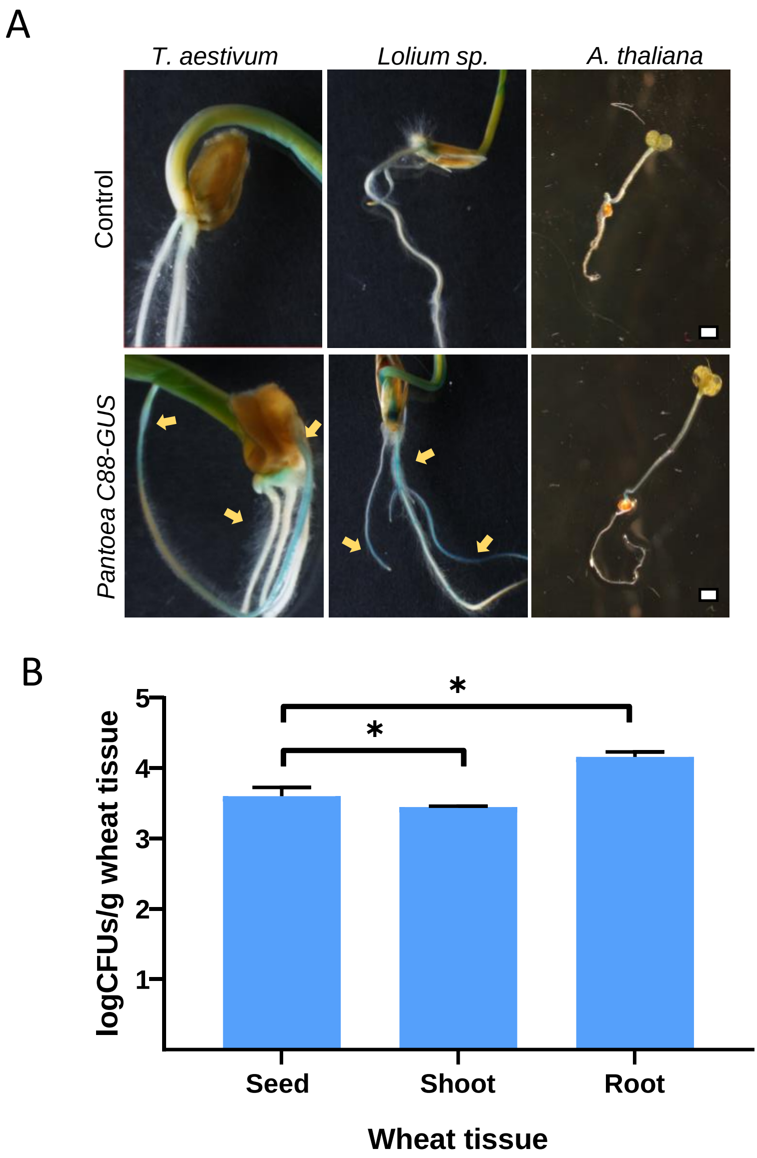


**Figure S4. *Pantoea* colonizes seeds and is transmitted to Poaceae seedlings.** A) Visualization of *Pantoea* colonization in wheat and *Lolium* seedlings using the chromogenic substrate X-gluc (yellow arrows). Bar 5 mm. B) Quantification of *Pantoea* populations in wheat tissues (seed, shoot, and root) by qPCR. Values represent the means of three independent experiments with three biological replicates per experiment. Error bars represent the standard deviation of the mean.

**Table S1.** Wheat samples used in this study that were harvested in Spain (_SP) from two different fields located in Asturias or Leon. Samples **in bold** were used in figures 3 and 4 (see Table S3). [a]Genotypes resulting from hybridizations and cultivated species: AA *T. monococcum* from *T. urartu*, AABB *T. turgidum* and *T. durum* from the hybridization with *Aegilops speltoides* (BB), AABBDD *T. aestivum* and *T.spelta*.

<table>
<tr><th>Sample</th><th>Specie</th><th>Location</th><th>Harvest Year</th><th>[a]Genome</th><th>Taming</th></tr>
<tr><td>Tmo_SP</td><td>T. monococcum</td><td>Asturias</td><td rowspan="3">2020</td><td>AA</td><td rowspan="2">Ancestral</td></tr>
<tr><td>Ttu_SP</td><td>T. turgidum</td><td rowspan="6">Leon</td><td rowspan="2">AABB</td></tr>
<tr><td>Tdu_SP</td><td>T. durum</td><td rowspan="5">Commercial</td></tr>
<tr><td>Tae_SP1</td><td rowspan="4">T. aestivum</td><td>2018</td><td rowspan="9">AABBDD</td></tr>
<tr><td>Tae_SP2</td><td>2019</td></tr>
<tr><td>Tae_SP4</td><td>2020</td></tr>
<tr><td>Tae_SP5</td><td>2021</td></tr>
<tr><td>Tpe_SP1</td><td rowspan="5">T. spelta</td><td rowspan="4">Asturias</td><td>2018</td><td rowspan="5">Ancestral</td></tr>
<tr><td>Tpe_SP2</td><td rowspan="2">2019</td></tr>
<tr><td>Tpe_SP3</td></tr>
<tr><td>Tpe_SP4</td><td rowspan="2">2020</td></tr>
<tr><td>Tpe_SP5</td><td>Leon</td></tr>
</table>

**Table S2.** Wheat seed samples used in this study that belong to the seedbank from the Institute of Plant Genetics and Crop Plant Research (IPK). The collection includes four different *Triticum* species and *Aegilops tauschii*, one of the three progenitors of the hexaploid wheat (85). [a]Genotypes resulting from hybridizations and cultivated species: DD *A. tauschii*, AA *T. monococcum* from *T. urartu*, AABB *T. dicoccom* and *T. durum* from the hybridization with *Aegilops speltoides* (BB), AABBDD *T. aestivum*.

<table>
<tr><th>Sample</th><th>IPK Code</th><th>Genus</th><th>Specie</th><th>Country</th><th>Location</th><th>Year</th><th>[a]Genome</th><th>Taming</th></tr>
<tr><td>Ata_RU</td><td>AE498</td><td rowspan="2">Aegilops</td><td rowspan="2">tauschii</td><td>Russia</td><td rowspan="2">Asia</td><td>1979</td><td rowspan="2">DD</td><td rowspan="6">Ancestral</td></tr>
<tr><td>Ata_UZ</td><td>AE246</td><td>Uzbekistan</td><td>1976</td></tr>
<tr><td>Tmo_BU</td><td>TRI1998</td><td rowspan="10">Triticum</td><td rowspan="2">monococcum</td><td>Bulgaria</td><td>Europe</td><td rowspan="7">1975</td><td rowspan="2">AA</td></tr>
<tr><td>Tmo_UN</td><td>TRI4323</td><td rowspan="2">Unknown</td><td rowspan="2">Unknown</td></tr>
<tr><td>Tdi_UN</td><td>TRI4168</td><td rowspan="2">dicoccum</td><td rowspan="5">AABB</td></tr>
<tr><td>Tdi_US</td><td>TRI445</td><td>United States</td><td rowspan="2">America</td></tr>
<tr><td>Tdu_AR</td><td>TRI448</td><td rowspan="3">durum</td><td>Argentina</td><td rowspan="6">Commercial</td></tr>
<tr><td>Tdu_AF</td><td>TRI4101</td><td>Afghanistan</td><td rowspan="2">Asia</td></tr>
<tr><td>Tdu_IR</td><td>TRI6263</td><td>Iran</td></tr>
<tr><td>Tae_GE</td><td>TKS</td><td rowspan="3">aestivum</td><td>Germany</td><td rowspan="2">Europe</td><td>2016</td><td rowspan="3">AABBDD</td></tr>
<tr><td>Tae_GR</td><td>TRI10765</td><td>Greece</td><td rowspan="2">1975</td></tr>
<tr><td>Tae_NZ</td><td>TRI7121</td><td>New Zealand</td><td>Oceania</td></tr>
</table>

**Table S3**. Plant and soil field samples used in this study. All samples were harvested in Leon, Spain in 2019.

| Sample | Species | Tissue | [a]Genome | Taming |
|---|---|---|---|---|
| Tdu_So | *T. durum* | soil | AABB | Commercial |
| Tdu_Ro | | root | | |
| Tdu_Sh | | shoot | | |
| Tdu_Pk | | spike | | |
| Tdu_Se | | seed | | |
| Ttu_So | *T. turgidum* | soil | | Ancestral |
| Ttu_Ro | | root | | |
| Ttu_Sh | | shoot | | |
| Ttu_Pk | | spike | | |
| Ttu_Se | | seed | | |
| Tpe_So | *T. spelta* | soil | AABBDD | |
| Tpe_Ro | | root | | |
| Tpe_Sh | | shoot | | |
| Tpe_Pk | | spike | | |
| Tpe_Se | | seed | | |
| Tae_So | *T. aestivum* | soil | | Commercial |
| Tae_Ro | | root | | |
| Tae_Sh | | shoot | | |
| Tae_Pk | | spike | | |
| Tae_Se | | seed | | |

**Table S4.** Primers used in this study. *with the corresponding Illumina adapters when required.

| Primer code | Sequence | Reference |
|---|---|---|
| 27_F | 5'-AGAGTTTGATCMTGGCTCAG-3' | (Lane, 1991) |
| 515_F* | 5’-GTGCCAGCMGCCGCGGTAA-3’ | (Marchesi et al., 1998) |
| 808_R* | 5’-GACTACHVGGGTATCTAATCC-3’ | (Caporaso et al., 2011) |
| 1522_R | 5'-AAGGAGGTGATCCANCCRCA-3' | (Weisburg et al., 1991) |
| GUS_F | 5'-ACTCATTACGGCAAAGTGTGGGTCA-3' | (Almasi et al., 2015) |
| GUS_R | 5'-TGGTGTAGAGCATTACGCTGCGAT-3' | (Almasi et al., 2015) |
| gusA_R | 5'-TTAGCTCACTCATTAGG-3' | This work |
| gusA_F | 5'-GTTCATAGAGATAACCT-3' | This work |
| gusA-FAM | 5'-AGCATCAGGGCGGCTATACGCC-3' | This work |
